# Phosphatase PP2A and microtubule pulling forces disassemble centrosomes during mitotic exit

**DOI:** 10.1101/182618

**Authors:** Stephen J. Enos, Martin Dressler, Beatriz Ferreira Gomes, Anthony A. Hyman, Jeffrey B. Woodruff

## Abstract

Centrosomes are major microtubule-nucleating organelles that facilitate chromosome segregation and cell division in metazoans. Centrosomes comprise centrioles that organize a micron-scale mass of protein called pericentriolar material (PCM) from which microtubules nucleate. During each cell cycle, PCM accumulates around centrioles through phosphorylation-mediated assembly of PCM scaffold proteins. During mitotic exit, PCM swiftly disassembles by an unknown mechanism. Here, we used *Caenorhabditis elegans* embryos to determine the mechanism and importance of PCM disassembly in dividing cells. We found that the phosphatase PP2A and its regulatory subunit SUR-6 (PP2A^SUR-6^), together with cortically directed microtubule pulling forces, actively disassemble PCM. In embryos depleted of these activities, ~25% of PCM persisted from one cell cycle into the next, resulting in cytokinetic furrow ingression errors, excessive centrosome accumulation, and embryonic death. Purified pp2A^SUR-6^ could dephosphorylate the major PCM scaffold protein SPD-5 *in vitro*. Our data suggest that PCM disassembly occurs through a combination of dephosphorylation of PCM components and catastrophic rupture of the PCM scaffold.

## INTRODUCTION

Centrosomes are micron-scale, membrane-less organelles that nucleate microtubule arrays. They are crucial for assembling and positioning the mitotic spindle, establishing membrane polarity, and asymmetric cell division. Centrosomes comprise a pair of nanometer-scale centrioles that organize a micron-scale mass of protein called pericentriolar material (PCM). PCM is required for proper centriole duplication (Dammermann et al., 2004; Loncarek et al., 2008) and determines the activity of centrosomes by serving as a concentration compartment for client proteins that nucleate microtubules (Conduit et al., 2015; Woodruff et al., 2014). During each cell cycle, the PCM assembles around centrioles in preparation for mitosis and then rapidly disassembles during mitotic exit, while the centrioles persist. Post-mitotic cells often lose their PCM and centrioles altogether, suggesting a tight coupling of centrosome assembly status to cellular differentiation. In fact, centrosome disassembly is essential for female gamete formation in several organisms (Borrego-Pinto et al., 2016; Mikeladze-Dvali et al., 2012; Pimenta-Marques et al., 2016) and terminal differentiation of heart tissue in mice (Zebrowski et al., 2015). However, the importance of centrosome disassembly for mitotically dividing cells is not known. Additionally, the mechanism driving PCM disassembly is not known in any context.

PCM forms through phosphorylation-regulated assembly of long coiled-coil proteins into micron-scale scaffolds. These scaffolds then recruit client proteins, such as microtubule-stabilizing enzymes and tubulin, which are needed for centrosome function (Conduit et al., 2015; Woodruff et al., 2014). It has been shown that Polo kinase phosphorylation of Cdk5Rap2, Centrosomin, and SPD-5 is essential for PCM assembly in vertebrates, flies, and *C. elegans*, respectively (Conduit et al., 2010; Lee and Rhee, 2011; Woodruff et al., 2015). Furthermore, Polo kinase phosphorylation of Centrosomin and SPD-5 directly enhances their assembly into supramolecular scaffolds in vitro (Conduit et al., 2014; Feng et al., 2017; Woodruff et al., 2015; Woodruff et al., 2017). These results imply that removal of these phosphate moieties is important for PCM disassembly, but this idea has yet to be tested.

In this study, we set out to determine how PCM disassembles during mitotic exit in *C. elegans* embryos. We demonstrate that depletion of the PP2A phosphatase or its regulatory subunit SUR-6 slows down disassembly of the SPD-5 scaffold. Eliminating microtubule-dependent pulling forces in addition to SUR-6 depletion inhibited SPD-5 scaffold disassembly even further. We further show that purified pp2A^sur-6^ complexes dephosphorylate SPD-5 in vitro and that shear forces are sufficient to disrupt PCM scaffolds in vitro. Our results suggest that *C. elegans* PCM disassembles through dephosphorylation and microtubule-driven rupture of the SPD-5 scaffold.

## RESULTS

### Depletion of PP2A^sur-6^ or microtubule-dependent pulling forces inhibits PCM disassembly in vivo

We used time-lapse microscopy to monitor PCM disassembly in *C.elegans* embryos expressing GFP-labeled SPD-5 (GFP::SPD-5), the main component of the PCM scaffold (Hamill et al., 2002; Woodruff et al., 2015; Woodruff et al., 2017) (Figure 1A and 1B). Confirming previous analysis (Decker et al., 2011; Woodruff et al., 2015), PCM localized around centrioles shortly after fertilization, then grew in size as the embryo progressed toward mitosis (Movie S1). After anaphase onset, PCM expanded rapidly and then disintegrated as material simultaneously transited toward the cell cortex and dissolved (Figure 1B). During disassembly, anterior PCM deformation was relatively isotropic, whereas posterior PCM deformation occurred primarily along the short axis of the embryo (Figure 1B). Quantification of PCM disassembly using semiautomated tracking and segmentation revealed that PCM mass peaks ~275 s after nuclear envelope breakdown (NEBD), corresponding to anaphase; PCM is no longer detectable 600 s after NEBD, corresponding to interphase of the next cell cycle (Figure 1C and 1D). We conclude that PCM disassembly is completed in < 6 min and involves rupture and dissolution of the SPD-5 scaffold.

**Figure 1.**
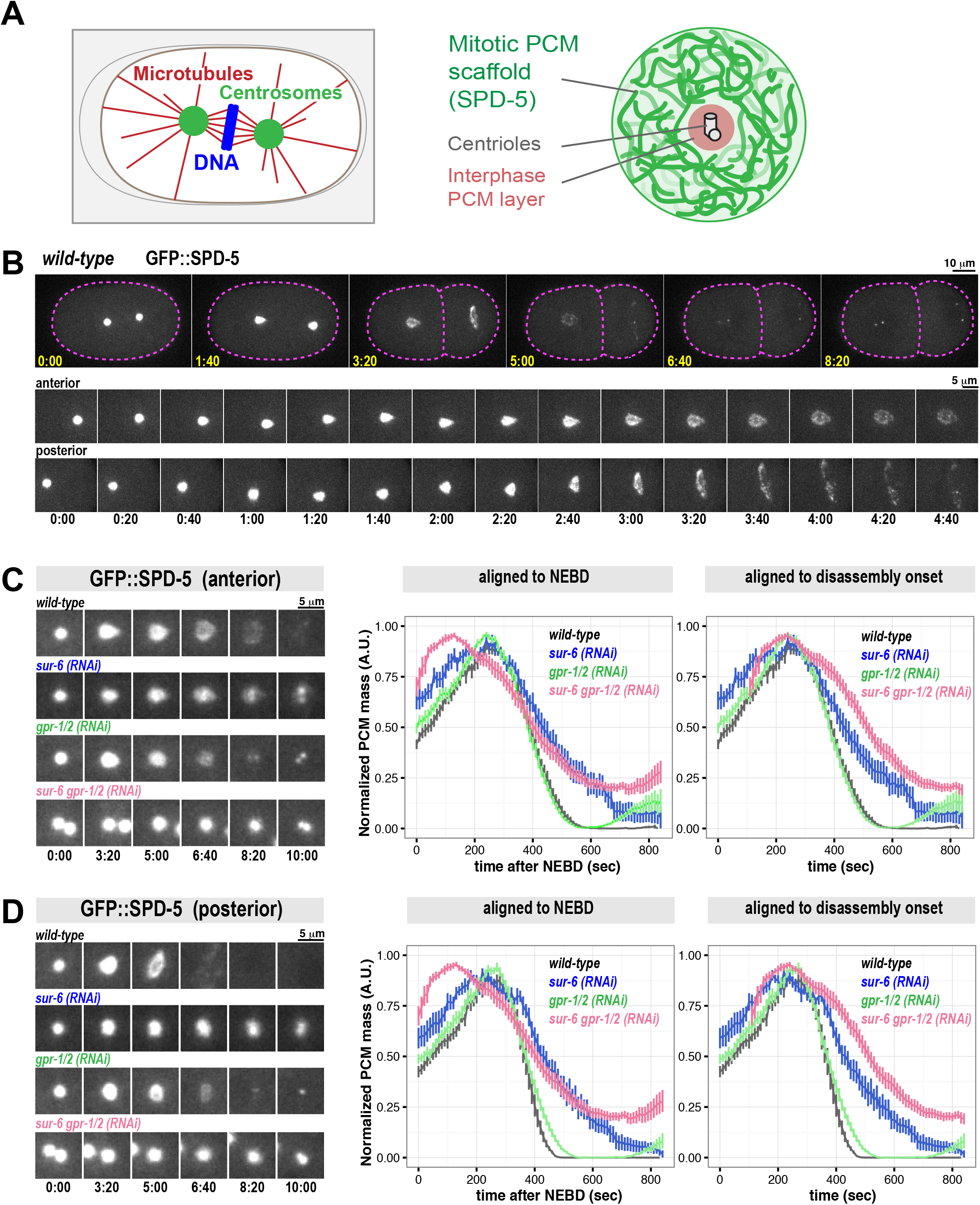
PCM disassembly is inhibited in *sur-6(RNAi)* and *gpr-1/2(RNAi)* embryos. A. Diagram of mitotic spindle (left) and centrosome organization (right). B. Confocal fluorescence images of *C. elegans* embryos expressing GFP::SPD-5, a marker for the PCM scaffold. During PCM disassembly in the 1-cell embryo, the anterior (left side) and posterior (right side) centrosomes display different morphologies. Cell outline is in magenta. Magnified images of the centrosomes are shown in the bottom panels. See also Movie S1. C. Measurement of anterior PCM disassembly in wild-type embryos and various mutant conditions (mean +/-SEM; n= 13 (wild-type), 9 *(sur-6(RNAi))*, 11 *(gpr-1/2 (RNAi)), 11(sur-6 gpr-1/2 (RNAi))*. Data are normalized. See Figure S1B for images of the embryos. D. Measurement of posterior PCM disassembly in wild-type embryos and various mutant conditions (mean +/-SEM; n= 13 (wild-type), 9 *(sur-6(RNAi))*, 11 *(gpr-1/2 (RNAi)), 11(sur-6 gpr-1/2 (RNAi))*. Data are normalized. See Figure S1B for images of the embryos.

PCM assembly is driven in part by PLK-1 (Polo-like Kinase) phosphorylation of SPD-5. In embryos, inhibition of PLK-1 or mutation of four PLK-1 target sites on SPD-5 prevents PCM growth (Woodruff et al., 2015; Wueseke et al., 2016). In vitro, PLK-1 phosphorylation of the same four sites accelerates assembly of SPD-5 into supramolecular scaffolds (Woodruff et al., 2015). To check whether dephosphorylation of these PLK-1 sites is critical for PCM disassembly, we performed a small-scale RNAi screen against known mitotic phosphatases. Only RNAi-mediated depletion of the PP2A phosphatase LET-92 inhibited PCM disassembly (Figure S1A; Movie S2). PP2A phosphatase localizes to centrosomes and connects to SPD-5 indirectly through the adapter proteins RSA-1 and RSA-2 (Schlaitz et al., 2007). Depletion of the catalytic subunit LET-92 causes pleiotropic effects that might indirectly affect PCM disassembly (Schlaitz et al., 2007). PP2A phosphatases function as holoenzymes comprising an invariant catalytic and structural subunit coupled to variable regulatory subunits that determine substrate specificity (Janssens and Goris, 2001). Depletion of the conserved B55a regulatory subunit (SUR-6 in *C. elegans* (Kao et al., 2004)) by RNAi prevented complete PCM disassembly without affecting spindle size or asymmetric cell division (Figure 1C and 1D; Figure S3A). *sur-6* depletion also reduced the speed of PCM disassembly (see Figure S1D for a comparison of disassembly rates). We conclude that PP2A coupled to SUR-6 (PP2A^SUR-6^) in part drives PCM disassembly.

Depletion of PP2A activity slowed down, but did not completely prevent, PCM disassembly, suggesting that additional mechanisms are required. Centrosomes are constantly under tension during anaphase due to pulling forces mediated by cortically-anchored dyneins that attach to and walk along astral microtubules emanating from PCM (Grill et al., 2001; Nguyen-Ngoc et al., 2007; Severson and Bowerman, 2003). In the *C. elegans* 1-cell embryo, pulling forces are xx-fold stronger in the posterior side compared to the anterior side (Grill et al., 2003). To test if microtubule-dependent pulling forces disassemble PCM, we knocked down the cortical dynein anchor GPR-1/2 by RNAi. In these mutant embryos, PCM still disassembled, albeit without the dramatic expansion in size seen in wild-type embryos (Figure 1C and 1D)(Severson and Bowerman, 2003). Quantification revealed a ~27% reduction in disassembly kinetics of the posterior centrosome (Figure 1D; Figure S1D). Thus, elimination of microtubule-pulling forces has a minor effect on disassembly of the posterior centrosome. Interestingly, PCM assembly was detectable much sooner in the subsequent cell cycle in *grp-1/2(RNAi)* embryos compared to wild-type embryos (Figure 1C and 1D). These results suggest that an active but barely detectable layer of PCM persists after mitotic exit in *grp-1/2(RNAi)* embryos; this layer which could prematurely seed PCM accumulation in the next cell cycle.

Combinatorial depletion of GPR-1/2 and SUR-6 resulted in a more severe PCM disassembly phenotype: PCM disassembly was ~2.5-fold slower than wild-type and ~25% of the original PCM mass persisted into the next cell cycle (Figure 1C and 1D; Figure S1D; Movie S3). We also noticed additional defects in the *sur-6 gpr-1/2(RNAi)* double mutant, such as mitotic spindle collapse and altered cell cycle progression (see next section). In particular, the time between NEBD and disassembly onset was shorter in the double mutant compared to the single mutant or wild-type embryos; thus, for comparison purposes, we display the PCM disassembly curves aligned by NEBD and by disassembly onset (Figure 1C and 1D). We conclude that pp2A^sur-6^ and microtubule pulling forces cooperate to disassemble PCM.

In early *C. elegans* embryos PCM grows until reaching a stereotyped upper limit (Decker et al., 2011). We wondered if PCM disassembly mechanisms help set this upper limit. Unexpectedly, we found that *gpr-1/2* depletion only slightly increased PCM mass in anaphase, while *sur-6* depletion actually decreased PCM mass (see Figure S1C for non-normalized data). PCM mass in anaphase in *sur-6 gpr-1/2(RNAi)* embryos was slightly lower than in wild-type embryos (Figure S1C). We conclude that the GPR-1/2 and SUR-6 disassembly pathways do not oppose PCM assembly prior to anaphase.

### PP2A^sur-6^ dephosphorylates a key PLK-1 site on SPD-5

SPD-5 assembly is accelerated by PLK-1 phosphorylation at four central serine residues within SPD-5 (S530, S627, S653, S658) (Woodruff et al., 2015). To determine if PP2A^sur-6^ dephosphorylates these residues, we generated a monoclonal antibody that specifically recognizes S530 only in the dephosphorylated state (Figure 2A). Western blot analysis showed that this antibody recognizes purified unphosphorylated SPD-5 in vitro. As expected, the antibody signal declined when SPD5 was phosphorylated by purified PLK-1 (Figure 2B). We refer to this antibody hereafter as “de-pS530”.

**Figure 2.**
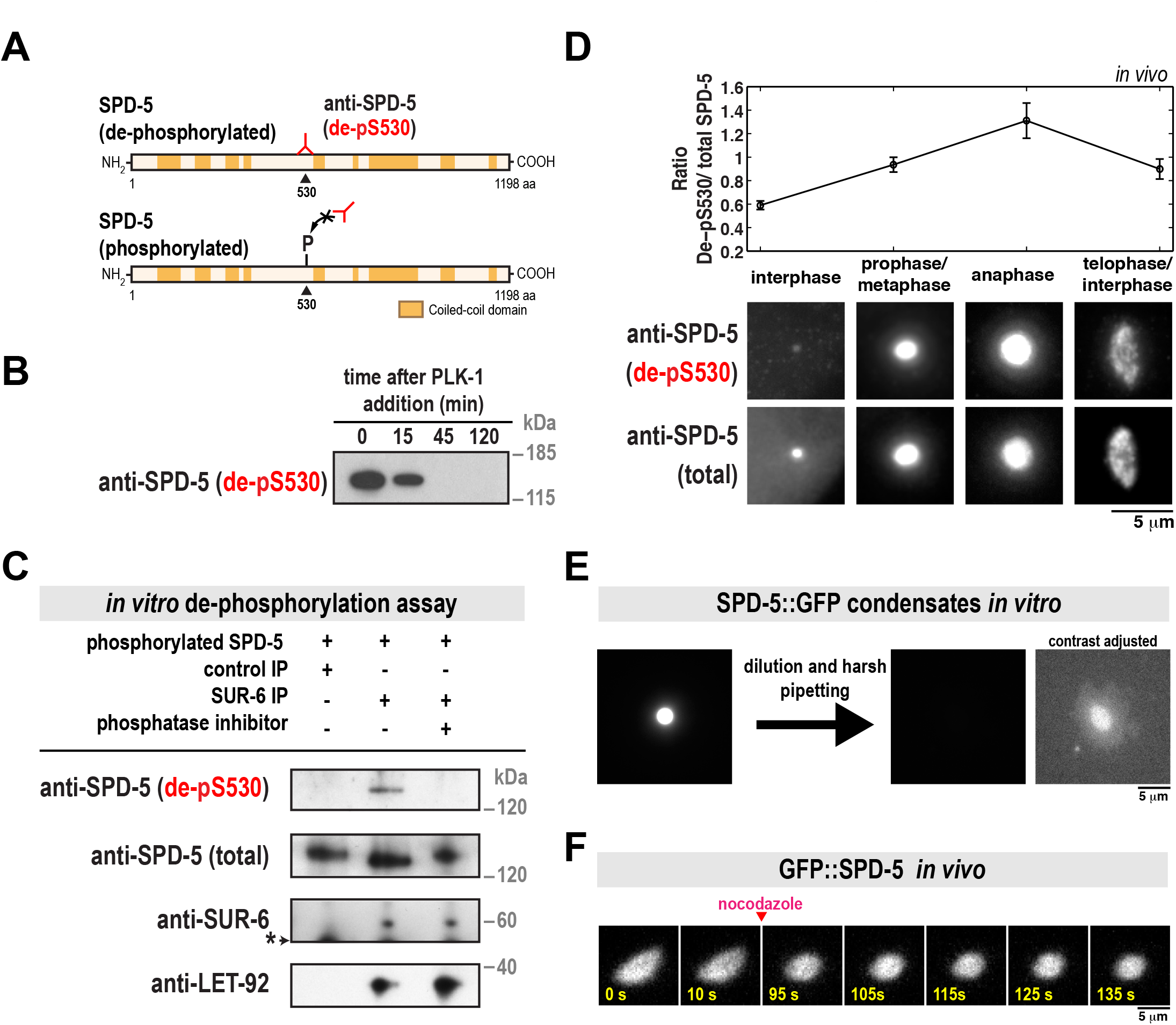
PP2A^sur-6^ dephosphorylates SPD-5 and shear stresses distort and dissolve SPD-5 assemblies in vitro. A. SPD-5 domain architecture and location of the serine 530 phospho-epitope. The de-pS530 antibody recognizes serine 530 only when dephosphorylated. B. 200 nM of purified SPD-5 was incubated with 200 nM PLK-1 + 0.2 mM ATP. The reaction was analyzed at various time points by western blot using the de-pS530 antibody. C. In vitro dephosphorylation assay. Control beads affixed to CDC-37 antibody or beads affixed to SUR-6 antibody were incubated in *C. elegans* embryo extract, then washed and resuspended in buffer. SPD-5 that was pre-phosphorylated in vitro by PLK-1 was then added and incubated for 90 min at 23°C and analyzed by western blot. Calyculin A was used as the phosphatase inhibitor (lane 3). The asterisk indicates a non-specific band. D. Dual color immunofluorescence images of *C. elegans* embryos using the de-pS530 antibody and a general polyclonal SPD-5 antibody (total SPD-5). Relative changes in SPD-5 dephosphorylation at different cell cycle stages were determined by measuring the ratio of the two antibody signals (de-pS530/total SPD-5)(mean +/-SEM; n = 19 interphase, 48 prophase/metaphase, 17 anaphase, and 33 telophase/interphase centrosomes). E. SPD-5 condensates were formed by incubating 500 nM SPD-5::GFP in 9% PEG-3350 for 27 min at 23°C (left), then diluted 1:10 into a 0% PEG solution and pipetted harshly (right). The two images on the right are the same, except the contrast has been increased in the far right image to show the disrupted SPD-5 condensate. F. Semi-permeable *perm-1(RNAi)* embryos expressing GFP::SPD-5 were treated with 20 ug/ml nocodazole after anaphase onset. PCM relaxes to a spherical shape after microtubules are depolymerized.

We next performed an in vitro dephosphorylation assay to test if PP2A^sur-6^ can directly dephosphorylate SPD-5 at S530. We affixed SUR-6 antibodies to beads to isolate pp2A^sur-6^ complexes from *C. elegans* embryo extracts. Western blot analysis confirmed that these complexes contained the regulatory subunit SUR-6 and the catalytic subunit LET-92 (Figure 2C). The pp2A^SUR-6^ beads were then resuspended in buffer that contained purified SPD-5 that had been pre-phosphorylated in vitro by PLK-1. As shown in Figure 2C, SPD-5 was dephosphorylated only in the presence of active PP2A^SUR-6^. de-pS530 signal was not present when control beads were used or if the PP2A inhibitor Calyculin A was included. These results suggest that pp2A^SUR-6^ drives PCM disassembly by dephosphorylating SPD-5.

We then performed dual color immunofluorescence with the de-pS530 antibody and a general SPD-5 antibody to assess relative changes in SPD-5 dephosphorylation during the cell cycle. The de-pS530 antibody localized to centrosomes in all cell cycle stages, corroborating our previous observation that a phospho-mutant version of SPD-5 (GFP::SPD-5^4A^) localizes to centrosomes when wild-type SPD-5 is present (Wueseke et al., 2016). Quantification of de-pS530 antibody signal showed that it was low during interphase, increased during metaphase, then peaked during anaphase, coincident with disassembly onset (Figure 2D). This result suggests that PCM disassembly is associated with dephosphorylation of SPD-5. Later in telophase and the subsequent interphase, de-pS530 signal decreased (Figure 2D). It is possible that dephosphorylated SPD-5 is weakly associated with the PCM and removed faster than phosphorylated SPD-5, as predicted by our previous model (Wueseke et al., 2016).

### Forces disassemble the SPD-5 scaffold in vitro and distort PCM in vivo

Our analysis of the *grp-1/2(RNAi)* mutant suggested that cortically directed pulling of microtubules assists pp2A^SUR-6^ in driving PCM disassembly. To test if force is sufficient to disassemble the PCM scaffold, we applied shear stress to SPD-5 assemblies in vitro. Purified SPD-5 forms PCM-like scaffolds that concentrate PCM client proteins like PLK-1, microtubule stabilizing enzymes, and tubulin. Depending on macromolecular crowding conditions, SPD-5 assembles either into dense spherical condensates or irregular networks (Woodruff et al., 2015; Woodruff et al., 2017). Application of shear force by harsh pipetting completely disassembled the less dense SPD-5 networks (Figure S2) and partially disassembled the denser SPD-5 condensates (Figure 2E). After harsh pipetting, some SPD-5 condensates lost their spherical morphology (Figure 2E). To validate these observations in vivo, we performed acute disruption of microtubule-dependent forces during anaphase by treating permeable embryos with 20 µg/ml nocodazole. After nocodazole application, the normally elongating PCM scaffold relaxed to a spherical shape (Figure 2F). Taken together, these results suggest that cortically directed forces are integral in rupturing and disassembling the PCM scaffold.

### PCM disassembly mutants display abnormal centrosome accumulation and improper cell divisions

PCM assembles and disassembles during each mitotic cell cycle in dividing metazoan cells. However, it is not clear why PCM must disassemble instead of persisting through each cell cycle like other organelles, such as mitochondria. We thus tested the impact of inhibiting PCM disassembly on embryo viability. *sur-6* null embryos *(sur-6(sv30))* or embryos treated with *gpr-1/2(RNAi)* for 24 hr at 23°C resulted in near 100% lethality (Gotta et al., 2003; Kao et al., 2004). When we reduced the strength of RNAi treatment, we observed a genetic interaction between *sur-6* and *gpr-1/2* (see Figure 3A for details). Under these conditions, lethality of F1 embryos was 0% in wild-type, 40% in *sur-6(RNAi)*, 5% in *gpr-1/2(RNAi)* and 60% in *sur-6 gpr-1/2(RNAi)* worms (Figure 3A). It is possible that this synthetic lethality results from inhibition of PCM disassembly. However, SUR-6 and GPR-1/2 are also known to regulate centriole duplication and spindle positioning, respectively (Gotta et al., 2003; Kao et al., 2004; Song et al., 2011). Disruption of these processes could contribute to the embryonic lethal phenotype. We therefore analyzed the earliest cell divisions in *C. elegans* embryos, which are known to be largely unaffected by single RNAi depletion of *sur-6* and *gpr-1/2* (Kao et al., 2004; Nguyen-Ngoc et al., 2007; Song et al., 2011).

**Figure 3.**
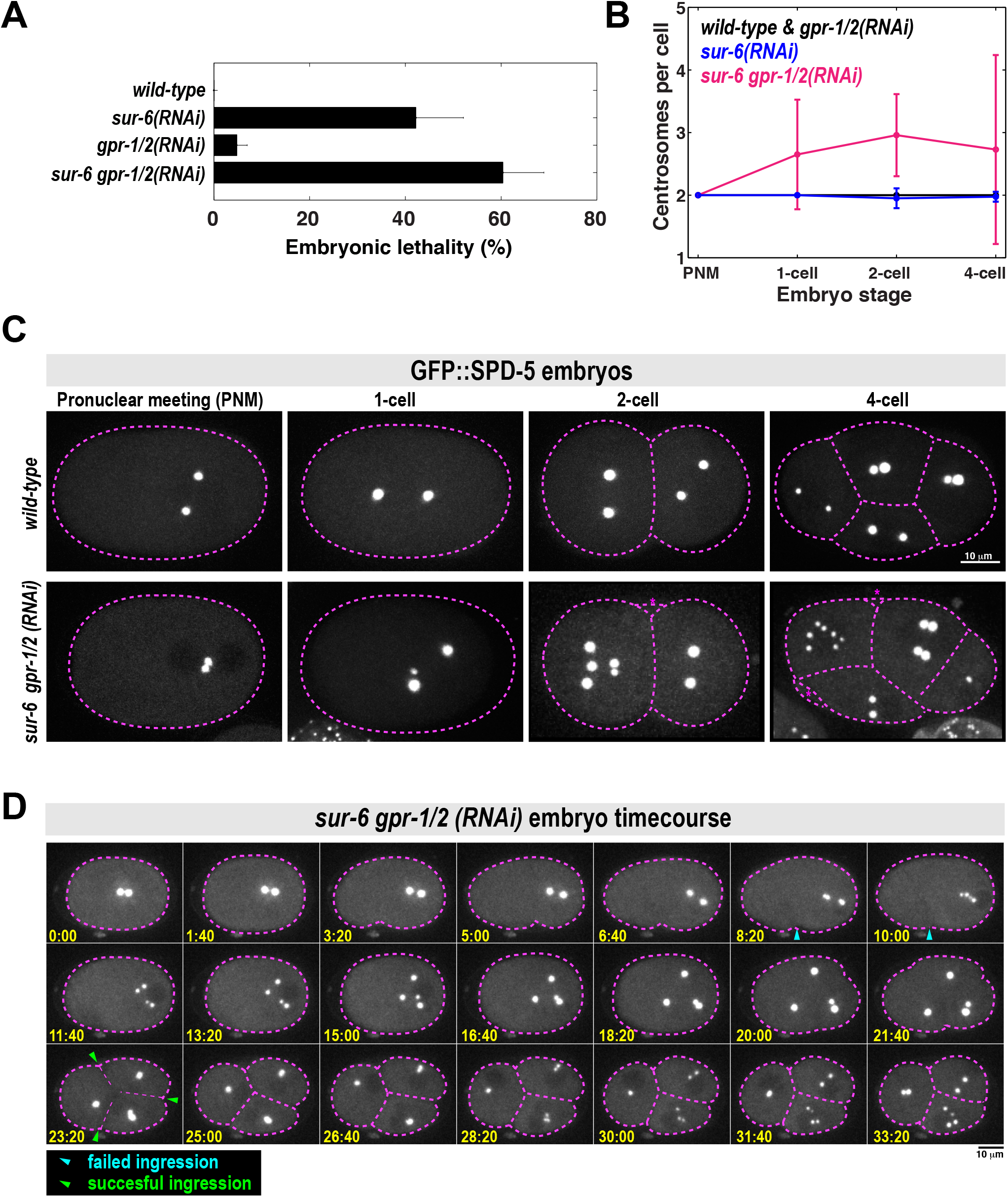
PCM disassembly mutants display abnormal centrosome numbers and cell divisions. A. Analysis of embryonic lethality in various partial RNAi conditions. For *sur-6(RNAi)*, L4 worms were grown on *sur-6* feeding plates for 16 hr at 23°C. For *gpr-1/2(RNAi)*, young adults were grown on *grp-1/2* feeding plates for 8 hr at 23°C. For *sur-6 gpr-1/2(RNAi)*, L4 worms were grown on *sur-6* feeding plates for 16 hr at 23°C, then transferred to *sur-6 gpr-1/2* feeding plates for an additional 8 hr at 23°C (n = 8 mothers per condition and >50 F1 embryos per mother). B. Number of centrosomes per cell in wild-type, *sur-6(RNAi), gpr-1/2(RNAi)* and *sur-6 gpr-1/2(RNAi)* embryos (mean +/-SD; n=10 embryos in each condition). All embryos contained only two centrosomes during pronuclear meeting (PNM), which occurs shortly after fertilization. 3-cell embryos were not counted, as they are transient in the wild-type condition and do not have visible centrosomes during that brief time. C. Representative images from (B). Cell outline is in magenta. Blebs were visible in the double mutant embryos, but were not counted as cells (magenta asterisks). D. Time-lapse imaging of a *sur-6 gpr-1/2(RNAi)* embryo expressing GFP::SPD-5. Blue arrowheads indicate abortive cytokinetic furrow ingression. Green arrowheads indicate successful furrow ingression. See also Movie S3.

We noticed severe mutant phenotypes in *sur-6 gpr-1/2(RNAi)* embryos that were not present in the single mutants. For example, metaphase spindles were noticeably shorter in the *sur-6 gpr-1/2(RNAi)* embryos compared to wild-type embryos and the single mutants (Figure S3A). Furthermore, wild-type and single mutant embryos contained only two centrosomes per cell, whereas *sur-6 gpr-1/2(RNAi)* embryos often contained >2 centrosomes; sometimes up to 7 centrosomes per cell could be seen by the 4-cell stage (Figure 3B and 3C).

We then followed embryo development with time-lapse microscopy to understand how excessive centrosome numbers arise in *sur-6 gpr-1/2(RNAi)* embryos (Figure 3D; Movie S3). All double mutant embryos contained only two centrosomes after fertilization, indicating that failed meiotic divisions were not responsible for the centrosome accumulation phenotype (Figure 3C and 3D; see pronuclear meeting stage). At the end of the first cell cycle, centrosomes separated, but cytokinetic furrow ingression failed. These centrosomes maintained PCM from the previous cell cycle but still managed to split into two new centrosomes and retain their spherical shape. They then accumulated PCM and formed new spindles, indicating cell cycle progression into the next mitosis.

Thus, we conclude that extra centrosomes accumulate due to failed cytokinesis combined with a normal centrosome duplication cycle. We sometimes saw odd numbers of centrosomes per cell due to incomplete centrosome splitting (Figure 3C and S3B). Because these phenotypes appear only in the double mutant, they likely arise from the genetic interaction between *sur-6* and *gpr-1/2*. Since we have shown that SUR-6 and GPR-1/2 cooperate during PCM disassembly, we propose that the mutant phenotypes are a consequence of failed PCM disassembly. However, we cannot discount the possibility that SUR-6 and GPR-1/2 cooperate in an unknown manner, independent of their effects on PCM disassembly, to regulate spindle assembly or cytokinesis.

## DISCUSSION

In this study we have shown that the PP2A phosphatase and cortically directed pulling forces are required for disassembly of PCM, the outer layer of centrosomes responsible for nucleating microtubules. Our results lead us to propose the following model for PCM assembly, maturation, and disassembly in *C. elegans* embryos. Prior to mitosis, PCM forms through self-assembly of the scaffold protein SPD-5 into micron-scale spherical condensates that then concentrate PCM client proteins, such as tubulin, needed for microtubule aster nucleation. SPD-5 scaffold formation is accelerated by the nucleator SPD-2 and PLK-1 phosphorylation of SPD-5. During mitotic exit, PP2A (LET-92 in *C. elegans)* coupled to its regulatory subunit B55α (SUR-6 in *C. elegans)* drives PCM disassembly by dephosphorylating the scaffold protein SPD-5, thereby opposing PLK-1. Simultaneously, microtubules emanating from PCM attach to the cortex and exert outward pulling forces that rupture the SPD-5 scaffold.

How does dephosphorylation of SPD-5 promote PCM disassembly? So far, we have not observed disassembly of SPD-5 condensates in the presence of phosphatase in vitro (unpublished data). This result suggests that once the SPD-5 scaffold is formed, dephosphorylation cannot destabilize it. This is possible considering that PLK-1 phosphorylation is not strictly required for SPD-5 assembly but rather affects the rate of assembly in vitro (Wueseke et al., 2016). Furthermore, the mature SPD-5 scaffold is incredibly stable and displays little to no turnover in vivo and in vitro, regardless of phosphorylation status (Laos et al., 2015; Woodruff et al., 2017; Wueseke et al., 2016). PP2A^SUR-6^ dephosphorylation of SPD-5 is also not sufficient to initiate PCM disassembly. For example, PCM disassembly occurs strictly in late anaphase, but our immunofluorescence experiments suggest that pp2A^SUR-6^ begins to dephosphorylate SPD-5 in metaphase. And, PCM does not disassemble after acute inhibition of PLK-1 in metaphase-arrested embryos (Wueseke et al., 2016). Lastly, cytoplasmic SPD-5 levels are relatively constant throughout the cell cycle (Wueseke et al., 2014), ruling out degradation of SPD-5 as a possible mechanism. Taken together, we propose that a PP2A^SUR-6^-independent mechanism initiates PCM disassembly after anaphase onset. PP2A^SUR-6^ dephosphorylation of SPD-5 then prevents SPD-5 reassembly after dissociating from the mature PCM. It is also likely that pp2A^SUR-6^ targets regulators of PCM assembly such as SPD-2, PLK-1, and Aurora A kinase; each of these proteins is activated in part by phosphorylation (Decker et al., 2011; Littlepage et al., 2002; Qian et al., 1998).

How can force-induced rupture drive PCM disassembly? Our data show that shear stress directly destabilizes SPD-5 assemblies in vitro. However, elimination of microtubule-pulling forces only slightly inhibits PCM disassembly in vivo. This could be a difference in force magnitude and the type of force applied, as pipetting (in vitro) would cause shear strain, while pulling (in vivo) would cause linear strain. It is also possible that PCM rupture increases the exposed surface of the PCM scaffold, making it more accessible to disassembly enzymes that would otherwise be excluded. This would not apply to PP2A, which concentrates at centrosomes (Schlaitz et al., 2007) and has a clear effect on PCM disassembly even when pulling forces are eliminated (Figure 1C). However, this principle could apply to as-of-yet unidentified disassembly mechanisms.

## Author Contributions

J.B.W., M.D., and A.A.H. conceived the project. J.B.W. wrote the manuscript. S.J.E. performed live-cell imaging and analyzed PCM disassembly in vivo. M.D. initially characterized the PCM disassembly mutants and performed the in vitro dephosphoryation assay. B.F.G. analyzed the in vivo data and performed in vitro disassembly experiments. J.B.W. performed all other experiments.

## Acknowledgements

We thank the Light Microscopy and Antibody facilities at the MPI-CBG; K. O’Connell for providing the SUR-6 antibody; Robert Haase for computational support; Andrea Zinke and Anne Schwager for help with worm maintenance. This project was funded by the Max Planck Society and the European Commission's 7th Framework Programme grant (FP7-HEALTH-2009-241548/MitoSys) and a MaxSynBio grant to A.H. J.B.W. was supported by an EMBO fellowship and MaxSynBio.

## METHODS

### Worm strain maintenance and RNA interference

*C. elegans* worm strains were grown on NGM plates at 16-23°C, following standard protocols (http://www.wormbook.org). We used one strain with the following genotype:

OD847: unc-119(ed9) III; ltSi202[pVV103/ p0D1021; Pspd-2::GFP::SPD-5 RNAi-resistant; cb-unc-119(+)]II

RNA interference was performed by feeding. For nocodazole treatment of embryos, L4 worms were grown on *perm-1(RNAi)* feeding plates at 20°C for 16-18 hr, then dissected in an open imaging chamber filled with osmotic support medium (Carvalho et al., 2011; Wueseke et al., 2016) and 20 µg/ml nocodazole (Sigma). For *sur-6* and *gpr-1/2 (RNAi)* treatment L4 worms were grown on their given plates at 23°C for 24-28 hr, then dissected and imaged following standard protocols.

### Imaging

For live embryo imaging, we used an inverted Olympus IX81 microscope with a Yokogawa spinning-disk confocal head (CSU-X1), a 60x 1.4 NA Plan Apochromat water objective, and an iXon EM + DU-897 BV back illuminated EMCCD camera (Andor). For analysis of PCM disassembly in vivo, we generated 36 X 0.5 µm Z-stacks every 10 s using 50 ms exposure and 8% laser intensity (4.5 mW; 488 nm laser). In vitro SPD-5 condensates and fixed embryos were visualized with an inverted Olympus IX71 microscope using 60x 1.42 NA or 100x 1.4 NA Plan Apochromat oil objectives, CoolSNAP HQ camera (Photometrics), and DeltaVision control unit (GE Healthcare).

### Assembly of SPD-5 condensates in vitro

SPD-5 condensates were formed by adding concentrated SPD-5::GFP to condensate buffer (25 mM HEPES, pH 7.4, 150 mM KCl) containing polyethylene glycol (molecular weight 3,350 Da) and fresh 0.5 mM DTT. See (Woodruff et al., 2015) for details on SPD-5 purification.

### In vitro kinase assay

For the experiment in Figure 2B, 200 nM SPD-5::GFP, 200 nM PLK-1::6xHis, and 0.1 mg/ml ovalbumin were incubated in kinase buffer (20 mM Tris, pH 7.4, 150 mM KCl, 10mM MgCl_2_, 0.2 mM ATP, 1 mM DTT) for 1 hr at 23° C.

### Centrosome tracking and quantification

For all centrosome disassembly measurements, acquired stacks were analyzed by a custom-made FIJI macro (see Figure S4).

#### Movie generation and alignment

First, Z stacks were collapsed into SUM projections at each time point and then combined to make time-lapse movies. Each movie was then split into two stacks to isolate the anterior and posterior centrosomes. The frame corresponding to nuclear envelope breakdown (NEBD) was identified, and only the frames after NEBD were processed.

#### Determination of background and threshold

On the first frame, a Gaussian blur was applied (sigma = 1) and the maximum intensity pixel was identified, which represents the center of the centrosome. A band-shaped region was created around the maximum intensity pixel with an inner circle that encompasses the largest extent of PCM signal (radius = 7 pixels) and an outer circle (radius = 10 pixels). Background mean intensity (meanbg) and standard deviation (stdevbg) were then calculated from the area between the two rings.

#### Segmentation and measurement of centrosomes

Centrosomes were segmented automatically by creating a region of interest (ROI) using the following threshold: mean_bg_ + 3*stdev_bg_. This threshold was applied to the remaining frames. The integrated signal intensity (a proxy for PCM mass) bounded by the ROI was calculated by: (mean_R0I_ - threshold)*area_R0I_. The data were normalized by dividing the integrated signal intensity of each frame by the highest value of integrated signal intensity of the stack. Since we were not able to separate the two centrosomes in the double RNAi strain, the image segmentation was done following the same method outlined above, except in two steps: the band-shaped region was created with a bigger radius and the integrated signal intensity measurements were divided by two.

### Immunofluorescence

Embryos were fixed in methanol and frozen in liquid N2, as previously described (Hamill et al., 2002), then stained with 1:2,000 anti-SPD-5 (de-pS530; mouse; BX23 clone) and 1:5,000 anti-SPD-5 (total; rabbit; 758 clone). 1:400 goat anti-rabbit-alexa488 and 1:400 goat anti-mouse-alexa594 (Life Technologies) were used as secondaries.

### In vitro dephosphorylation assay and western blotting

Worms were harvested in IP buffer (1× PBS plus 100 mM KCl, 1 mM EGTA, 1 mM MgCl_2_, 1% CHAPS, and 1× Complete Protease Inhibitor cocktail (Roche)) and snap-frozen in liquid nitrogen. Frozen worm pellets were turned into powder using a Retsch MM301 mill. Worm lysate was prepared by resuspending worm powder in 1.5ml IP buffer per gram powder. Lysate was cleared by centrifuging for 10 min at 10,000 x g at 4°C. The cleared lysate was again centrifuged for 10 min at 16,000 x g at 4°C. Immunoprecipitation was carried out using Dynabeads Protein G kit (Life Technologies) and anti-SUR-6 and anti-CDC-37 antibodies. Instead of eluting the protein after the immunoprecipitation, the buffer was exchanged for PP2A kinase buffer (40 mM Tris pH 8.4, 34 mM MgCl_2_, 4 mM EDTA, 2 mM DTT, 0.05 mg/ml BSA). SUR-6/LET-92 bound to beads was used to dephosphorylate recombinant SPD-5 phosphorylated in vitro using recombinant PLK-1 (see In vitro kinase assay; the reaction was passed over a Ni-NTA column to remove PLK-1::6xHis). Dephosphorylation was carried out at room temperature for 1.5 hr in PP2A kinase buffer. Aliquots of the reactions were separated on 4-12% NuPAGE gradient gels (Life Technologies). Proteins were transferred onto nitrocellulose membrane using an iBlot device (Life Technologies). Membranes were blocked using 3% BSA in TBST containing 0.1% Tween-20 and probed using antibodies against total SPD-5 (1:5000; 758 clone, in-house), SPD-5 which is not phosphorylated at S530 (1:1000; BX23 clone, in-house), SUR-6 (1:500, gift from K. O’Connell) and PP2A catalytic subunit (1:2,000; BD Biosciences). Secondary antibodies were HRP-conjugated goat anti-rabbit and goat anti-mouse (Bio-Rad, 1:30,000). Detection was carried out using SuperSignal ECL reagent (Biorad).

## SUPPLEMENTAL FIGURE LEGENDS

**Figure S1.**
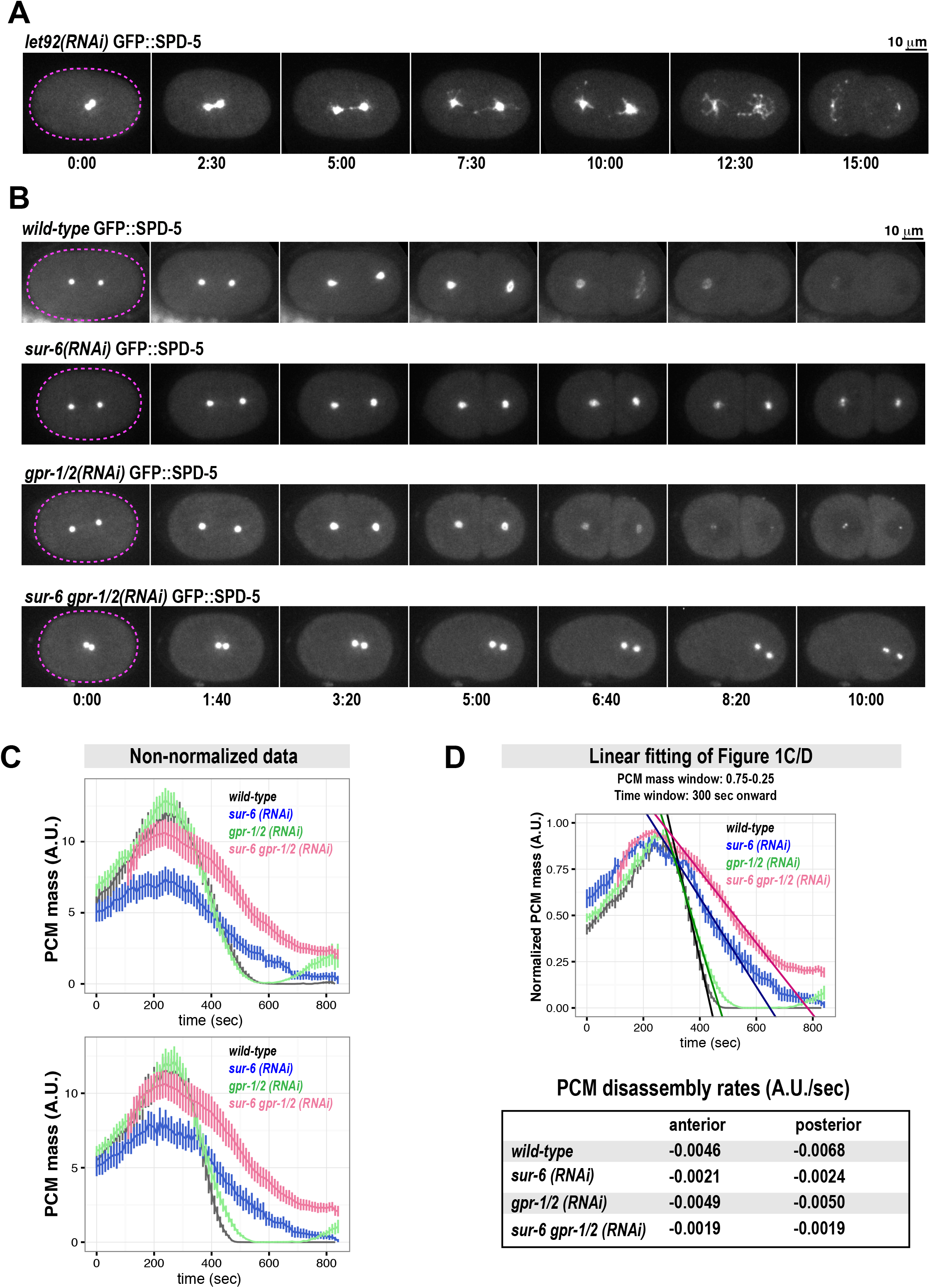
PCM disassembly defects in various mutant conditions. A. Confocal images of an embryo expressing GFP::SPD-5 after *let-92* was depleted by RNAi (24 hr at 23° C). See also Movie S2. B. Images of the entire embryos from the analysis in Figure 1C and 1D. C. Non-normalized quantification of PCM mass from Figure 1C and 1D. Curves are aligned by peak PCM mass. D. PCM disassembly rates calculated by fitting the linear region of the curves in Figure 1C and 1D (linear region: window between 0.75 and 0.25 PCM mass during disassembly, corresponding to 300 s post-NEBD). An example of the fitting is shown in the plot on the right.

**Figure S2.**
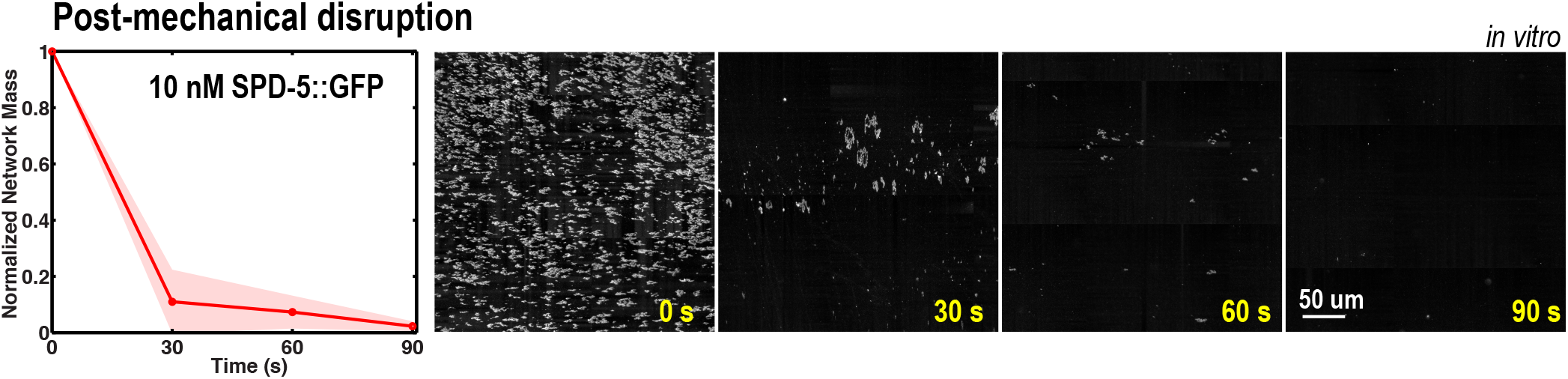
Harsh pipetting dissolves SPD-5 networks in vitro. After 1 hr at 23° C, 10 nM SPD-5::GFP assembled into micron-scale networks. The sample was then pipetted 10 times and network mass (integrated fluorescence) was calculated at different time points as described in (Woodruff et al., 2015). The red line represents the mean, the shaded area represents the 95% confidence interval (n = 9 experiments).

**Figure S3.**
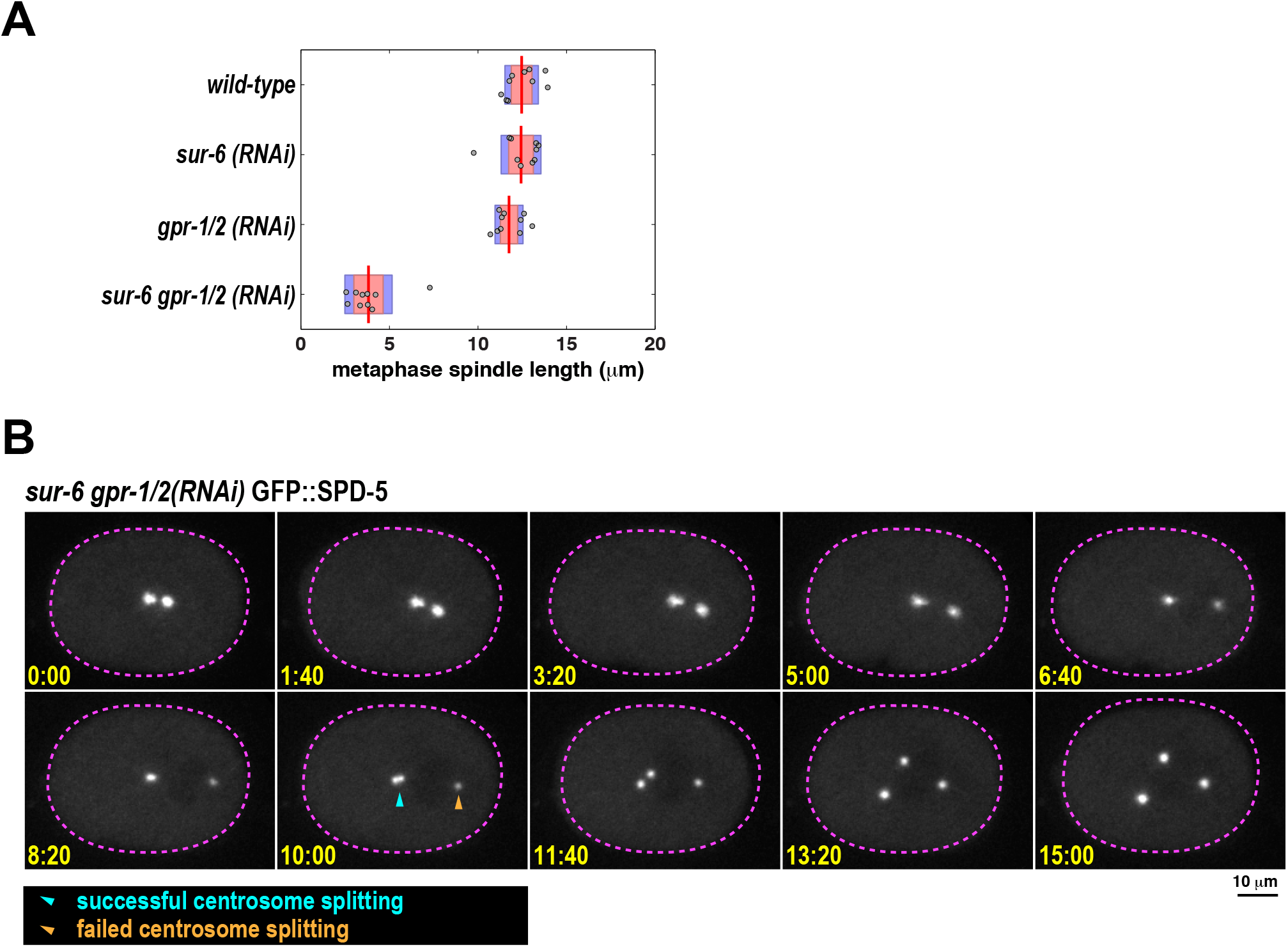
Analysis of metaphase spindle length and centrosome splitting in PCM disassembly mutants. A. Metaphase spindle length in 1-cell embryos from Figure 1C and 1D. Plot shows means (red bars), 95% confidence intervals (red shaded areas), and S.D. (blue shaded areas; n = 10 spindles per condition). B. Example of a sur-6 gpr-1/2(RNAi) embryo where one centrosome fails to split (orange arrowhead) after onset of PCM disassembly.

**Figure S4.**
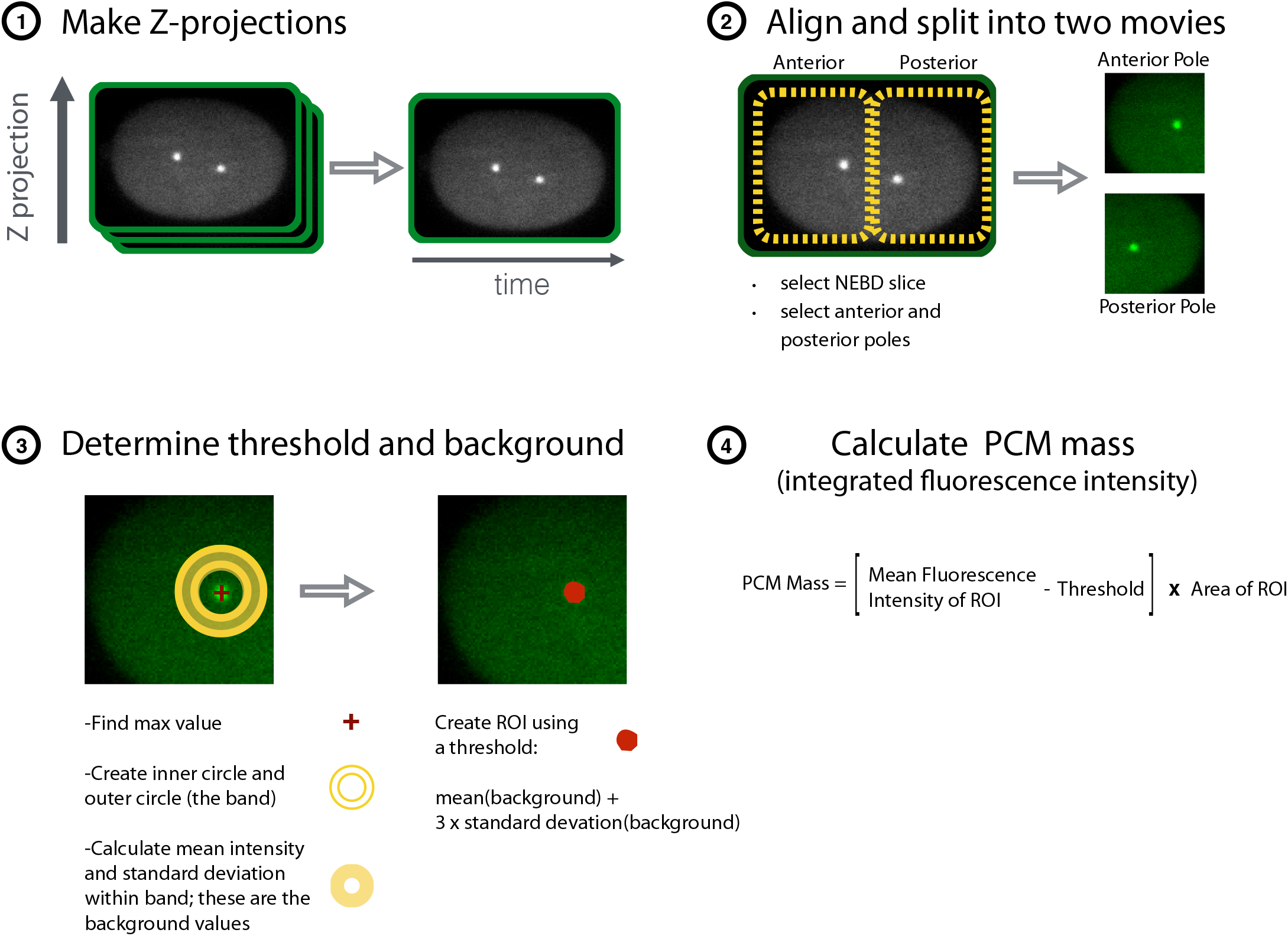
Outline of PCM disassembly quantification. See methods for a detailed explanation.

## Supplemental movies

Movie S1. PCM assembly and disassembly in a one-cell wild-type embryo expressing GFP::SPD-5.

Movie S2. PCM disassembly in a *let-92(RNAi)* embryo expressing GFP::SPD-5.

Movie S3. PCM disassembly in a *sur-6 gpr-1/2(RNAi)* embryo expressing GFP::SPD-5.

## REFERENCES

Borrego-Pinto, J., Somogyi, K., Karreman, M. A., König, J., Müller-Reichert, T., Bettencourt-Dias, M., Gönczy, P., Schwab, Y. and Lénárt, P. (2016). Distinct mechanisms eliminate mother and daughter centrioles in meiosis of starfish oocytes. The Journal of Cell Biology 212, 815–827.

Carvalho, A., Olson, S. K., Gutierrez, E., Zhang, K., Noble, L. B., Zanin, E., Desai, A., Groisman, A. and Oegema, K. (2011). Acute drug treatment in the early *C. elegans* embryo. PLoS ONE 6, e24656.

Conduit, P. T., Brunk, K., Dobbelaere, J., Dix, C. I., Lucas, E. P. and Raff, J. W. (2010). Centrioles Regulate Centrosome Size by Controlling the Rate of Cnn Incorporation into the PCM. Current Biology 20, 2178–2186.

Conduit, P. T., Feng, Z., Richens, J. H., Baumbach, J., Wainman, A., Bakshi, S. D., Dobbelaere, J., Johnson, S., Lea, S. M. and Raff, J. W. (2014). The centrosome-specific phosphorylation of Cnn by Polo/Plk1 drives Cnn scaffold assembly and centrosome maturation. Developmental Cell 28, 659–669.

Conduit, P. T., Wainman, A. and Raff, J. W. (2015). Centrosome function and assembly in animal cells. Nat Rev Mol Cell Biol 16, 611–624.

Dammermann A., Müller-Reichert T., Pelletier L., Habermann B., Desai A. and Oegema K. (2004). Centriole assembly requires both centriolar and pericentriolar material proteins. Developmental Cell 7, 815–829.

Decker, M., Jaensch, S., Pozniakovsky, A., Zinke, A., O'Connell, K. F., Zachariae, W., Myers, E. and Hyman, A. A. (2011). Limiting Amounts of Centrosome Material Set Centrosome Size in *C. elegans* Embryos. Current Biology 21, 1259–1267.

Feng, Z., Caballe, A., Wainman, A., Johnson, S., Haensele, A. F. M., Cottee, M. A., Conduit, P. T., Lea, S. M. and Raff, J. W. (2017). Structural Basis for Mitotic Centrosome Assembly in Flies. Cell 169, 1078-1089.e13.

Gotta, M., Dong, Y., Peterson, Y. K., Lanier, S. M. and Ahringer, J. (2003). Asymmetrically distributed *C. elegans* homologs of AGS3/PINS control spindle position in the early embryo. Curr. Biol. 13, 1029–1037.

Grill, S. W., Gönczy, P., Stelzer, E. H. and Hyman, A. A. (2001). Polarity controls forces governing asymmetric spindle positioning in the Caenorhabditis elegans embryo. Nature 409, 630–633.

Grill, S. W., Howard, J., Schäffer, E., Stelzer, E. H. K. and Hyman, A. A. (2003). The distribution of active force generators controls mitotic spindle position. Science 301, 518–521.

Hamill, D. R., Severson, A. F., Carter, J. C. and Bowerman, B. (2002). Centrosome Maturation and Mitotic Spindle Assembly in *C. elegans* Require SPD-5, a Protein with Multiple Coiled-Coil Domains. Developmental Cell 3, 673–684.

Janssens V. and Goris J. (2001). Protein phosphatase 2A: a highly regulated family of serine/threonine phosphatases implicated in cell growth and signalling. Biochem. J. 353, 417–439.

Kao, G., Tuck, S., Baillie, D. and Sundaram, M. V. (2004). *C. elegans* SUR-6/PR55 cooperates with LET-92/protein phosphatase 2A and promotes Raf activity independently of inhibitory Akt phosphorylation sites. Development 131, 755–765.

Laos, T., Cabral, G. and Dammermann, A. (2015). Isotropic incorporation of SPD-5 underlies centrosome assembly in *C. elegans*. Curr. Biol. 25, r648–9.

Lee K. and Rhee K. (2011). PLK1 phosphorylation of pericentrin initiates centrosome maturation at the onset of mitosis. 195, 9.

Littlepage, L. E., Wu, H., Andresson, T., Deanehan, J. K., Amundadottir, L. T. and Ruderman, J. V. (2002). Identification of phosphorylated residues that affect the activity of the mitotic kinase Aurora-A. Proc. Natl. Acad. Sci. U.S.A. 99, 15440–15445.

Loncarek, J., Hergert, P., Magidson, V. and Khodjakov, A. (2008). Control of daughter centriole formation by the pericentriolar material. Nature Cell Biology 10, 322–328.

Mikeladze-Dvali, T., von Tobel, L., Strnad, P., Knott, G., Leonhardt, H., Schermelleh, L. and Gönczy, P. (2012). Analysis of centriole elimination during *C. elegans* oogenesis. Development 139, 1670–1679.

Nguyen-Ngoc, T., Afshar, K. and Gönczy, P. (2007). Coupling of cortical dynein and G alpha proteins mediates spindle positioning in Caenorhabditis elegans. Nature Cell Biology 9, 1294–1302.

Pimenta-Marques, A., Bento, I., Lopes, C. A. M., Duarte, P., Jana, S. C. and Bettencourt-Dias, M. (2016). A mechanism for the elimination of the female gamete centrosome in Drosophila melanogaster. Science 353, aaf4866–aaf4866.

Qian, Y. W., Erikson, E. and Maller, J. L. (1998). Purification and cloning of a protein kinase that phosphorylates and activates the polo-like kinase Plx1. Science 282, 1701–1704.

Schlaitz, A.-L., Srayko, M., Dammermann, A., Quintin, S., Wielsch, N., MacLeod, I., de Robillard, Q., Zinke, A., Yates, J. R., III, Müller-Reichert, T., et al. (2007). The *C. elegans* RSA Complex Localizes Protein Phosphatase 2A to Centrosomes and Regulates Mitotic Spindle Assembly. Cell 128, 115–127.

Severson A. F. and Bowerman B. (2003). Myosin and the PAR proteins polarize microfilament-dependent forces that shape and position mitotic spindles in Caenorhabditis elegans. J Cell Biol 161, 21–26.

Song, M. H., Liu, Y., Anderson, D. E., Jahng, W. J. and O'Connell, K. F. (2011). Protein phosphatase 2A-SUR-6/B55 regulates centriole duplication in *C. elegans* by controlling the levels of centriole assembly factors. Developmental Cell 20, 563571.

Woodruff, J. B., Ferreira Gomes, B., Widlund, P. O., Mahamid, J., Honigmann, A. and Hyman, A. A. (2017). The Centrosome Is a Selective Condensate that Nucleates Microtubules by Concentrating Tubulin. Cell 169, 1066-1077.e10.

Woodruff, J. B., Wueseke, O. and Hyman, A. A. (2014). Pericentriolar material structure and dynamics. Philos. Trans. R. Soc. Lond., B, Biol. Sci. 369.

Woodruff, J. B., Wueseke, O., Viscardi, V., Mahamid, J., Ochoa, S. D., Bunkenborg, J., Widlund, P. O., Pozniakovsky, A., Zanin, E., Bahmanyar, S., et al. (2015). Centrosomes. Regulated assembly of a supramolecular centrosome scaffold in vitro. Science 348, 808–812.

Wueseke, O., Bunkenborg, J., Hein, M. Y., Zinke, A., Viscardi, V., Woodruff, J. B., Oegema, K., Mann, M., Andersen, J. S. and Hyman, A. A. (2014). The Caenorhabditis elegans pericentriolar material components SPD-2 and SPD-5 are monomeric in the cytoplasm before incorporation into the PCM matrix. Molecular Biology of the Cell 25, 2984–2992.

Wueseke, O., Zwicker, D., Schwager, A., Wong, Y. L., Oegema, K., Jülicher, F., Hyman, A. A. and Woodruff, J. B. (2016). Polo-like kinase phosphorylation determines Caenorhabditis elegans centrosome size and density by biasing SPD-5 toward an assembly-competent conformation. Biology Open 5, 1431–1440.

Zebrowski, D. C., Vergarajauregui, S., Wu, C.-C., Piatkowski, T., Becker, R., Leone, M., Hirth, S., Ricciardi, F., Falk, N., Giessl, A., et al. (2015). Developmental alterations in centrosome integrity contribute to the post-mitotic state of mammalian cardiomyocytes. Elife 4, 461.

